# A multi-state model of chemoresistance to characterize phenotypic dynamics in breast cancer

**DOI:** 10.1101/207605

**Authors:** Grant R. Howard, Kaitlyn E. Johnson, Areli Rodriguez Ayala, Thomas E. Yankeelov, Amy Brock

**Affiliations:** Department of Biomedical Engineering, The University of Texas at Austin, Austin, Texas 78712 USA; Department of Chemical Engineering, The University of Texas at Austin, Austin, Texas 78712 USA; Center for Computational Oncology, Institute for Computational Engineering Sciences, The University of Texas at Austin, Austin, Texas 78712 USA; Livestrong Cancer Institutes, Dell Medical School, The University of Texas at Austin, Austin, Texas USA; Diagnostic Medicine, Dell Medical School, The University of Texas at Austin, Austin, Texas USA

**Keywords:** (5–10): tumor, mathematical modeling, chemoresistance, doxorubicin, MCF-7, heterogeneity, dose-response, time-resolved microscopy

## Abstract

The development of resistance to chemotherapy is a major cause of treatment failure in breast cancer. Although several molecular mechanisms of chemotherapeutic resistance are well studied, a quantitative understanding of the dynamics of resistant subpopulations within a heterogeneous tumor cell population remains elusive. While mathematical models describing the dynamics of heterogeneous cancer cell populations have been proposed, few have been experimentally validated due to the complex nature of resistance that limits the ability of a single phenotypic marker to sufficiently isolate drug resistant subpopulations. In this work, we address this problem with a combined experimental and modeling system that uses drug sensitivity data to reveal the composition of multiple subpopulations differing in their level of drug resistance. We calibrate time-resolved dose-response data to three mathematical models to interrogate the models’ ability to capture the dynamics of drug. All three models demonstrated an increase in population level resistance following drug exposure. The candidate models were compared by Akaike information criterion and the model selection criteria identified a multi-state model incorporating the role of population heterogeneity and cellular plasticity. To validate the ability of this model to identify the composition of subpopulations, we mixed wild-type MCF-7 and MCF-7/ADR resistant cells at various proportions and evaluated the corresponding model output. Our blinded two-state model was able to estimate the proportions of cell subtypes, with the measured proportions falling within the 95 percent confidence intervals on the parameter estimations and at an R-squared value of 0.986. To the best of our knowledge, this contribution represents the first work to combine experimental time-resolved drug sensitivity data with a mathematical model of resistance development.

## Introduction

We aim to investigate how the therapeutic sensitivity of a breast cancer cell population changes over time following exposure to a pulse of the drug. We hypothesize that intratumoral heterogeneity and cellular plasticity play a direct role in the progression of resistance. This hypothesis is based on previous work demonstrating that exposure to chemotherapy induces gene expression changes, metabolic state transitions, and increased drug resistance in subsets of cancer cells^1–11^. In this work, we address this problem by using mathematical modeling to determine the frequencies of cells in different drug sensitivity states over time.

Approximately 30 percent of women diagnosed with early-stage breast cancer develop resistance and ultimately progress to metastastic breast cancer^12^. Doxorubicin is a standard-of-care cytotoxic agent in breast cancer treatment; however, the average time to develop resistance to doxorubicin is only 6 to 10 months^12^. Thus, it is critical to develop a model to describe the conditions and dynamics associated with the onset of chemotherapeutic resistance *in vitro* to begin to improve *in vivo* treatment regimens. We and others have demonstrated evidence of cellular plasticity and adaptability in response to chemotherapeutic treatment^1–6^. In a recent study, it was revealed that melanoma cells exhibit heterogeneity in their metabolic state, with cells utilizing different amounts of oxidative phosphorylation and aerobic glycolysis^6^. Here functional heterogeneity played a direct role in drug resistance as inhibition of aerobic glycolysis led to an increase in sensitivity to treatment^6^. The ability of individual cells to transition from a drug-sensitive to drug-resistant state has been observed in HL60 leukemia cells following chemotherapy exposure. Pisco *et al.* demonstrated that a subpopulation of cells increases expression of the ABC-transporter protein MDR1 in response to a chemotherapeutic pulse, leading to increased drug efflux and increased chemoresistance in those cells^1^. These experimental results focus on specific drug resistance phenotypes that emerge in cell subpopulations following treatment. However, because of the vast complexity of resistance mechanisms, it is difficult to identify a single molecular marker of drug resistance that encompasses all drug resistant cells^13,14^.

Mathematical descriptions of the dynamics of drug resistance may play a critical role in the development of strategies to combat this challenge ^13,15,16^. Theoretical models have been proposed that incorporate heterogeneous subpopulations in predicting and optimizing treatment response^17–23^; however, these models have not been fully validated with experimental cell population data *in vitro* or *in vivo.* While these approaches that incorporate the heterogeneity of resistant and sensitive subpopulations are promising, they remain mostly theoretical in nature^16^. Strategies such as optimal control theory^18,19^ (treatment aimed at maintaining the optimal composition of cell subpopulations), have yet to be implemented in patients because of lack of experimental validation. Knowledge of the predicted subpopulations proposed in these models is essential in order to progress from theory to implementation.

Although resistance to chemotherapy is a major cause of failure in breast cancer, we do not currently have a model describing the development of resistance in the context of a dynamic heterogeneous cancer cell population. On the other hand, experimental evidence concerning the variety of biological mechanisms of drug resistance is largely derived from static biological observations^24,25^. Many *in vitro* models of resistance have utilized chemoresistant cell lines established by gradually increasing the dose of a chemotherapeutic agent over time. In some cases, these experimental models have a median lethal dose (LD50) up to 14 times higher than the original cell line^25^, which may fail to capture the dynamics of the clinical onset of chemoresistance. The morphology, drug-sensitivity, and gene expression profiles of resistant cell lines have been analyzed in an attempt to tie these static observations to mechanisms of resistance^25^. While these studies are essential for discovering potential mechanisms, they do not aid in our understanding of the dynamics of resistance in response to brief doses of chemotherapeutic drug. The aim of this contribution is to examine the changes in drug resistance over time of a breast cancer cell population to add to the understanding of time-dependency mechanisms that lead to the resistant phenotype.

In this contribution, we calibrated experimental drug sensitivity data to multiple dynamic population models to test the hypothesis that there is a time-dependent population response to a chemotherapy treatment, and that this response is best described by models that incorporate heterogeneity and cellular plasticity. Plasticity is captured as changes in the frequency of cell state subpopulations. We combine the functional relevance of experimentally observed drug resistance data with various mathematical models to reveal the dynamic proportions of cells in subpopulations defined by their drug resistance. To validate that our modeling system was able to identify the composition of a cell population, we applied the model to known mixtures of reference cell populations of different resistance levels. To the best of our knowledge, this is the first effort that experimentally captures the time course of response to drug perturbation and provides a means of identifying the composition of cells in distinct drug sensitivity states.

## Materials and Methods

### Data acquisition Methods

#### Cell Culture

MCF-7 human breast cancer cells were obtained from ATCC and maintained in MEM (Thermo Fischer) supplemented with 10% fetal bovine serum (Gibco) and 1% Penicillin-Streptomycin (Gibco). MCF-7/ADR human breast cancer cells were generated in the laboratory of Dr. Kenneth Cowan (National Cancer Institute) and were a kind gift of Dr. Robert Clarke (Georgetown University)^26^. They were maintained in MEM (Gibco) supplemented with 10% fetal bovine serum (Gibco), 1% Penicillin-Streptomycin (Gibco), and 500 nM doxorubicin (Sigma-Aldrich). A subline of MCF-7 breast cancer cell line was engineered to constitutively express EGFP with a nuclear localization signal (EGFP-NLS). Genomic integration of the EGFP expression cassette was accomplished utilizing the Sleeping Beauty transposon system (doi: 10.1038/ng.343). The EGFP-NLS sequence was ordered as a gBlock from IDT and cloned into the optimized sleeping beauty transfer vector pSBbi-Neo. pSBbi-Neo was a gift from Eric Kowarz (Addgene plasmid #60525) ^27^. To mediate genomic integration, this two-plasmid system consisting of the transfer vector containing the EGFP-NLS sequence and the pCMV(CAT)T7-SB100 plasmid containing the Sleeping Beauty transposase was co-transfected into the MCF-7 population utilizing Lipofectamine 2000. mCMV(CAT)T7-SB100 was a gift from Zsuzsanna Izsvak (Addgene plasmid # 34879) ^28^. GFP^+^ cells were collected by fluorescence activated cell sorting. This MCF-7-EGFPNLS1 cell line is maintained in MEM (Gibco) supplemented with 10% fetal bovine serum and 200 μg/mL G418 (Caisson Labs).

#### Time Resolved Resistance Measurement

MCF-7 cells were plated at 6600 cells/cm^3^ and cultured for two days in growth media. The media was then exchanged for growth media containing 500 nM doxorubicin. After 24 hours, the doxorubicin media was washed out and replaced with growth media to end the drug pulse. Cells were passaged and counted weekly and drug sensitivity assays were performed weekly, as described below. Cell number counts at each week were used to determine the average per capita growth rate per day of the recovering cell population (**Fig 1a**).

**Figure 1.**
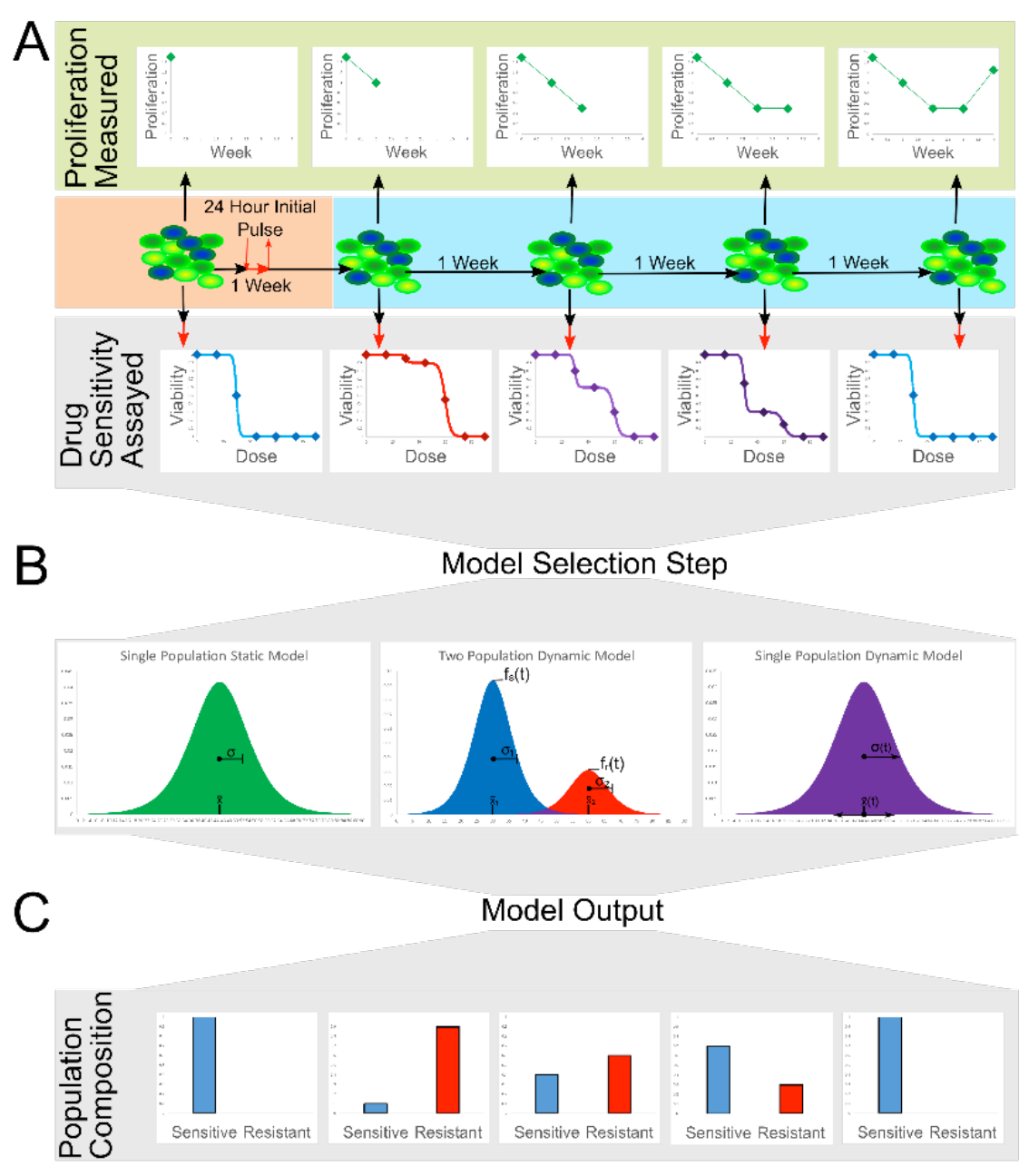
Experimental and Modeling Workflow. **a.** MCF-7 breast cancer cells are treated with an initial pulse of doxorubicin (500 nM) for 24 hours. Following drug removal, the sensitivity of the cell population is assessed weekly by measuring LD50. Proliferation rates are also measured each week. **b.** Using the dose response curve at each time point, multiple models are tested to determine the best means of capturing the dynamic cell population response. Model selection statistics indicate that a multipopulation model of at least two subpopulations is the best model. **c.** The model output of population structure at each time point along with overall proliferation rates can be used to estimate the number of resistant and sensitive cells over time following drug perturbation.

#### Drug Sensitivity Assay

300,000 cells were plated into a 12-well plate in growth media. After 2 days of culture, media was exchanged for growth media containing doxorubicin at a range of concentrations (0, 4, 14, 24, 36, 48, 60, 72, 84, 96, 120, and 144 μM). Twenty-four hours after this dosing, the cells (including supernatant media) were collected via trypsinization, pelleted, and resuspended in 20 μL of media. Live and dead cells were identified with acridine orange and propidium iodide (ViaStain AOPI Staining Solution, Nexcelom Bioscience) and quantified with a Nexcelom Cellometer VBA. The ratios of live to dead cells were used to determine the viability at each dose of doxorubicin (**Fig 1a**).

#### Cell Mixtures for Model Validation

MCF-7-EGFPNLS1 and MCF-7-ADR cells were counted, mixed at desired ratios (1:0, 3:1, 1:1, 1:3, and 0:1), and plated in 12-well plates as described for drug sensitivity assays. For each defined mixture, a sample of the untreated sample was counted in the Nexcelom Cellometer VBA to determine the actual observed ratio, using the EGFP fluorescence of the MCF-7-EGFP-NLS1 cell line as a marker.

### Data Analysis Methods

#### Calibration of Experimental Data to Multiple Structural Models of Dose-Response

The time-resolved dose response data was fit to three different models (**Box 1**). To perform an estimation of the parameters for all three models, a nonlinear least squares approach was implemented in MATLAB. Sigmoidal curves are chosen as the main structural model because they are often used to describe chemotherapy dose-response curves^29^ and contain physically identifiable parameters of a center and slope corresponding to the LD50 and reciprocal of the spread in the population distribution of drug-tolerance, respectively. The simplest model is a model of all data as a single homogenous cell population with a single static LD50 and slope. The single static population model equation is:

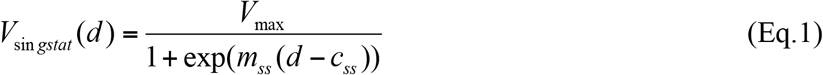

where *V* is the proportion of cells viable at the *d*, dose of doxorubicin in μM applied, *c*_*ss*_ is the LD50 of the entire population over all time, *m*_*ss*_ is slope at which the cells die as dose increases, which is the reciprocal of the standard deviation of the population, and *V*_*max*_ is the maximum viability of the cell population (as measured by the assay in absence of drug). This term is included to normalize for naturally occurring cell death independent of the effects of doxorubicin. The single static model represents the null hypothesis that the initial pulsed dose has no time-dependency in its effect on the cancer cell population.

##### Box 1.

###### Mathematical Models

The equations used for each of the three different structural models that were fit the time-resolved dose-response data. “Model Equation” gives the functional form of the equation, with (t) representing a parameter that was fit to the data set at each week. “Variables and Parameters” describes the variables used in terms of their physical meaning and their relation to the time-resolved data set.

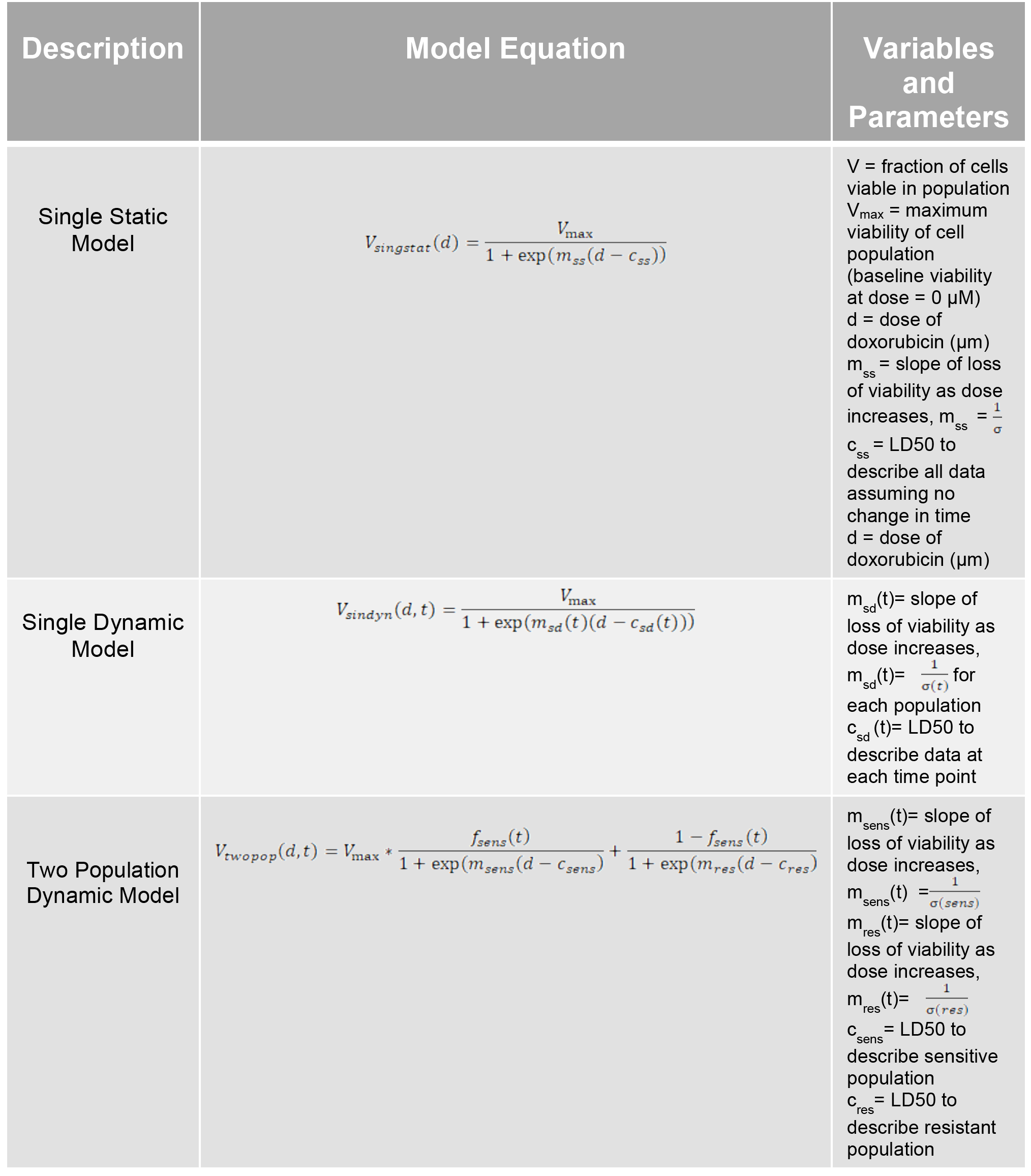

The second model is a model of all data as a single homogenous population whose population-level drug tolerance changes over time. The single dynamic population model equation is:

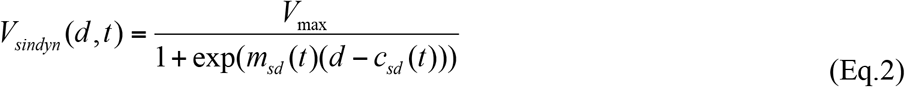

In this model, the *c*_*sd*_ and *m*_*sd*_ (LD50 and slope) parameters were allowed to float at each time point tested, leading to a 16-parameter model (slope and LD50 at each of the 8 weeks).

The third model is a model of a two-state system in which the fraction of cells in each state changes over time. The two-state dynamic population model equation is:

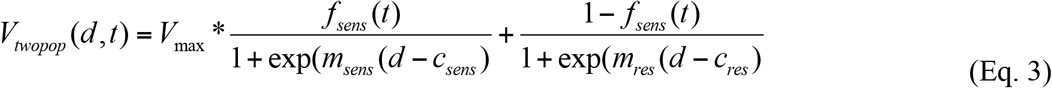

where each cell state is modeled as a subpopulation of cells whose LD50 is centered about a mean and slope (*c*_*sens*_, *m*_*sens*_ and *c*_*res*_, *m*_*res*_) which remained constant over time and whose *f*_*sens*_ and *f*_*res*_ (1- *f*_*sens*_) parameters were fit at each time point to best capture each week’s dose response data. The overall cell population viability (measured) is modeled as a direct sum of the viability response in each subpopulation. To fit the two-state model to data from multiple time points, the parameters of the sensitive and resistant slope and LD50 were forced to be kept constant over all time points, and the sensitive and resistant fraction parameters were allowed to float at each week, leading to a 12-parameter model (4 fixed parameters, 8 time dependent fraction parameters for each of the 8 weeks).

#### Statistical Analysis and Model Selection

For all three models, the confidence intervals on the parameter estimates were found using the bootstrapping method of replacement, with 500 boot-strapped simulated data sets. For each model, the mean-squared error and the Akaike Information Criterion (AIC) were calculated for stand-alone model statistics. The Akaike Information Criterion estimator (AIC value) was used for direct model comparison. The AIC value evaluates a model based on goodness of fit and penalizes for the complexity of the model using the number of parameters, with a lower AIC value indicating a better model. These evaluation criteria are used to decide on the most appropriate model to describe the dynamic dose response (**Fig 1b**).

#### Model Validation

To validate our modeling approach, we tested our models’ ability to reveal known mixtures of wild-type MCF-7 cells with MCF-7/ADR resistant cell line. The same fitting algorithm is used to fit the two-population dynamic model, except in this case we allowed each known group of mixture replicates to have their own fractional parameter, and maintained that the LD50 and centers of the two populations remain constant over all data. The bootstrapping method of replacement, again with 500 simulated data sets, was used to assess the confidence intervals around our parameter estimates of fraction of sensitive and resistant cells within a sample.

## Results

### Cancer Cell Population Exhibits Time-Dependent Response to Pulse Treatment

To determine whether the MCF-7 population level resistance varies in time, we fit the dynamic data to both the static and dynamic single population model, as shown in **Figure 2a** & **b**. In **Figure 2a**, all combined measured data is shown alongside the single static population model curve, for both the untreated controls (black) and the pulse-treated cell populations (green). The LD50 to describe the resistance of these populations is 37.0 +/- 3.5 μM for the untreated, and 50.4 +/- 2.4 μM for the treated population. In **Figure 2b**, we show the output of the single dynamic model parameters of the estimated LD50 values with the 95% confidence intervals in their estimation over all 8 weeks. The Akaike Information Criterion estimator (AIC value) is used for direct model comparison. The lower AIC value of the single dynamic model (-2015.6) compared to the single static model (-2004.4) indicates the time-resolved dose-response data are better described by the single dynamic model that allows the LD50 value of the population to change at each time point (**Table 1**). The pattern in parameter estimations of the LD50 values in the dynamic model (**Fig 2b**) corroborate this statistical analysis, showing a significant peak in the estimated LD50 at week 2 at 67.2 +/- 10.0 μM, followed by a slow return towards baseline, reach 46.4 +/- 5.1 μM at 8 weeks.

**Figure 2.**
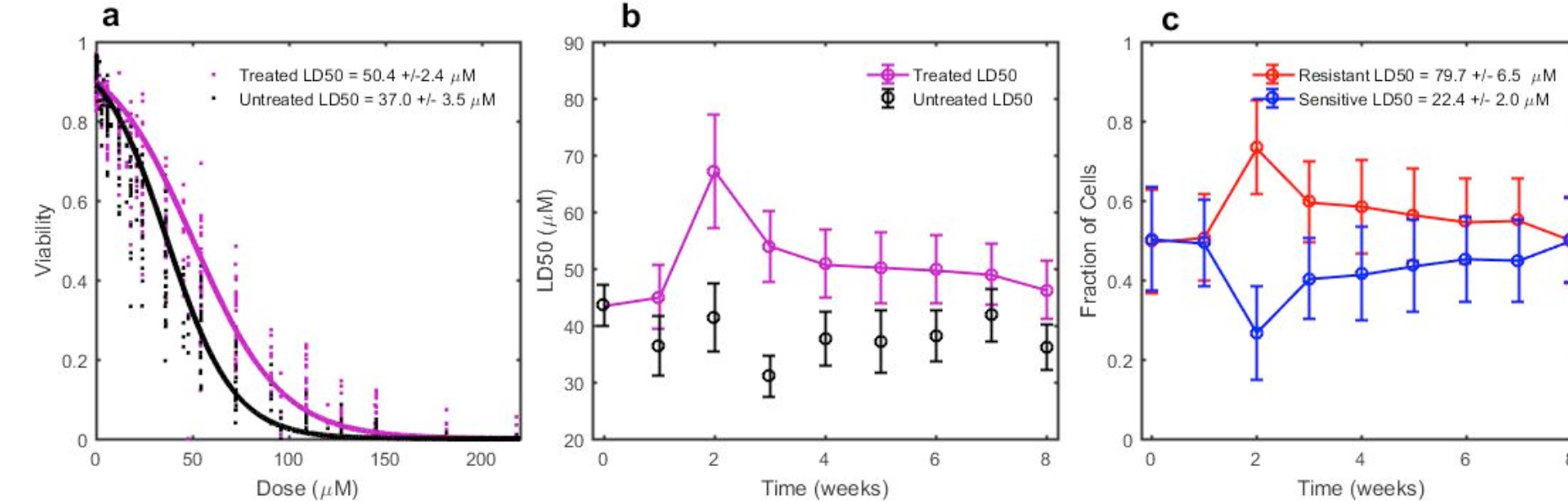
Time-resolved Drug Sensitivity Data Fit to Multiple Models. **a.** The static single population model demonstrates that the average resistance over 8 weeks significantly increases following exposure to pulsed treatment of doxorubicin **b.** A dynamic single population model displays the LD50 of the populations over time, demonstrating that the increase is time-dependent, peaking at 2 weeks after treatment, followed by a return towards baseline resistance levels. **c.** A two population dynamic model displays the model-estimated proportions of a population with LD50= 797.7 +/- 6.5 μM (resistant) and a population with LD50= 22.4 +/- 2.0 μM (sensitive). The relative proportions of these two populations change over time, yielding the observed LD50 for the overall population. Parameter fits at each time point are connected by a line for visual aid.

**Table 1.**
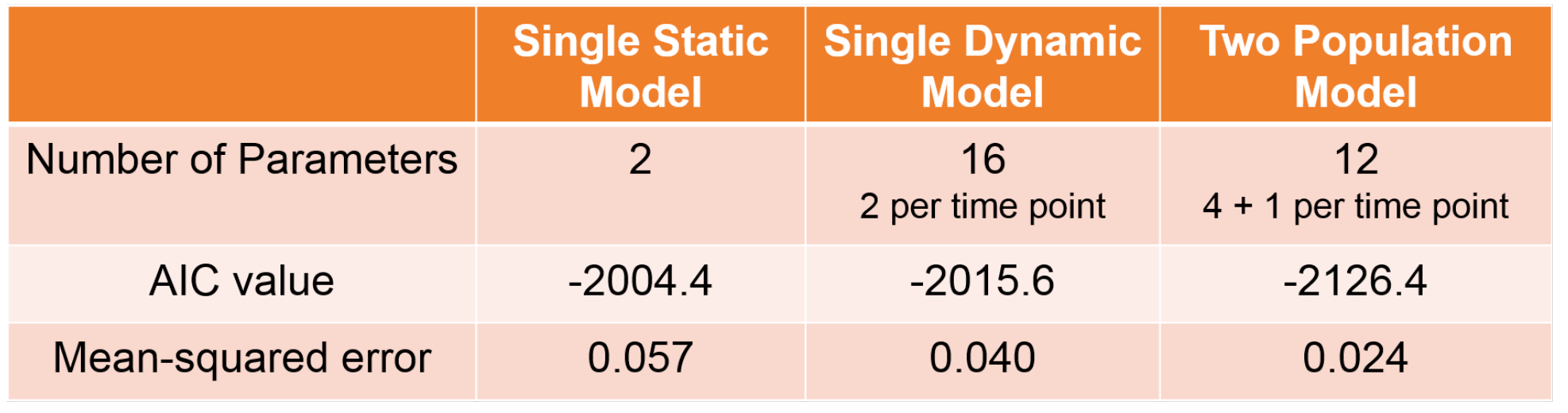
Model fit and model selection statistics indicate two population model improves fit and minimizes parameters. The dynamic two population model has the lowest AIC value and the lowest mean-squared error, indicating that the two-population model is both significantly improved in terms of model selection and in terms of error minimization than the single dynamic population model or the single static population model.

|  | Single Static Model | Single Dynamic Model | Two Population Model |
| --- | --- | --- | --- |
| Number of Parameters | 2 | 16<br>2 per time point | 12<br>4 + 1 per time point |
| AIC value | -2004.4 | -2015.6 | -2126.4 |
| Mean-squared error | 0.057 | 0.040 | 0.024 |

### Incorporating Heterogeneity via Drug Sensitivity States Improves Description of Response

To determine whether the dynamic drug resistance could be explained by a model of two subpopulations, we fit the data to the two-state model and output the resulting sensitive and resistant fraction parameter estimations, with their 95% confidence intervals, in Figure 2c. The two-state model describes two subpopulations of a cell population with distinct LD50 values whose relative frequencies are able to change in time after initial chemotherapy exposure, but the population centers remain constant. The two-state dynamic model outputs a resistant subpopulation with an LD50 of 79.7 μM +/- 6.5 μM and a sensitive subpopulation with an LD50 of 22.4 +/- 2.0 μM (**Fig 2c**). We again use model selection to demonstrate that the two-state model is an improved means of describing this time-resolved drug response in cell population dynamics. Support for selection of this model is indicated from the lower AIC value (-2126.4) of the two-population model compared with the single dynamic model (-2004.4) (**Table 1**). To illustrate the improvement in fit to the data, **Figure 3a** displays the dose-response data at 2 and 8 weeks with the two-state model curve for those weeks overlaid. This is supported by the lower mean-squared error value of 0.024 of the two-state model over the single dynamic model with a mean-squared error value of 0.040, despite it having less parameters than the single dynamic population model (**Table 1**).

**Figure 3.**
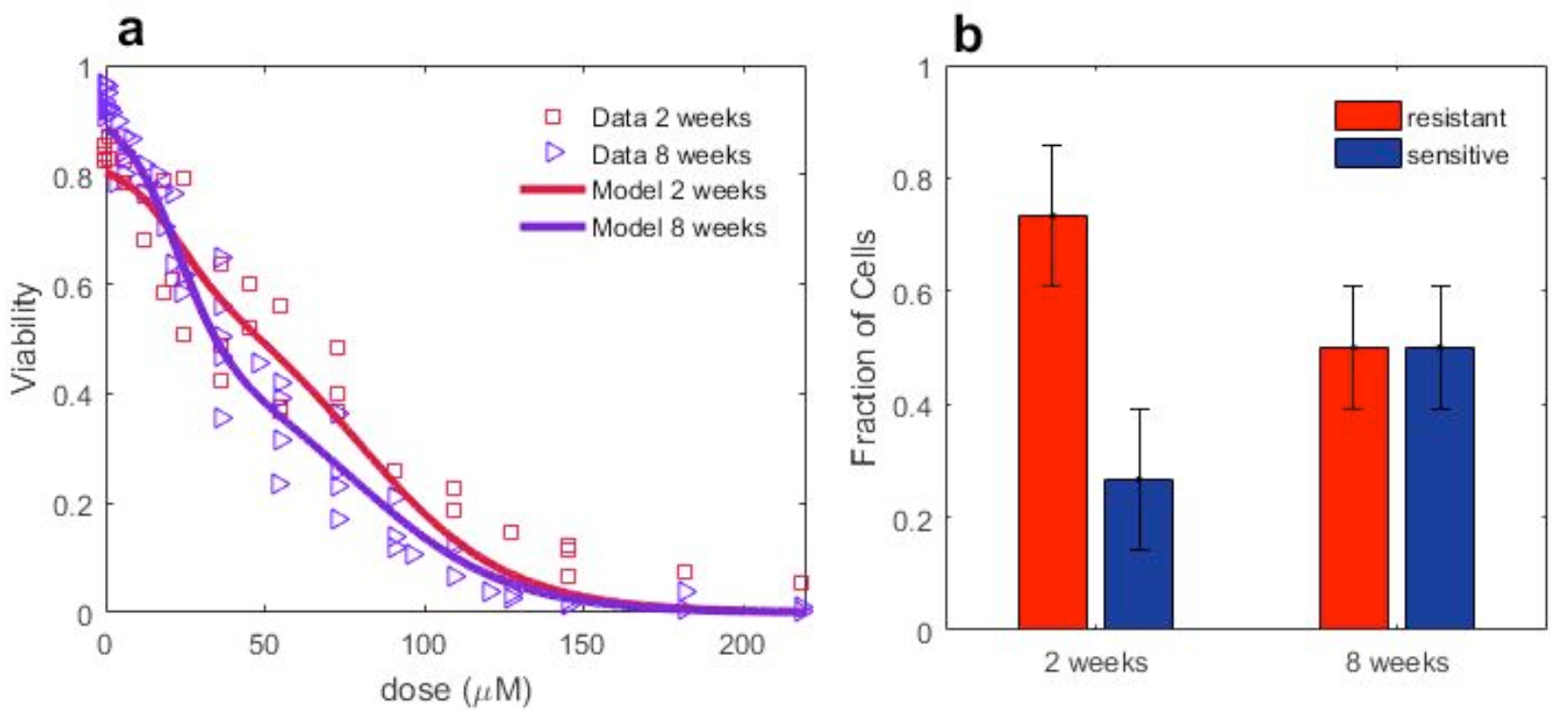
Example of Fit of Drug Sensitivity to the Two-population Model. **a.** Best fit of the two-population model to the dose response data at two and eight weeks post-treatment demonstrates the ability of the model to capture the differences in subpopulation levels at the different time points. **b.** Two population model output of parameter values of resistant and sensitive fractions at 2 and 8 weeks.

### Subpopulation Levels of Resistant Cells Transiently Increase

The key result for the treatment of doxorubicin on the MCF-7 cell line is that the proportion of cells in the resistant state is consistently higher than baseline from weeks 2 to 5 after the initial drug pulse, followed by return towards the initial resistant and sensitive subpopulation levels (**Fig 1c**). Changes in the proportions of cell subtypes can be combined with weekly measured per capita growth rate per day (birth minus death) (**Fig 4a**) to obtain estimates of the number of resistant and sensitive cells over time in a treated population (**Fig 4b**).

**Figure 4.**
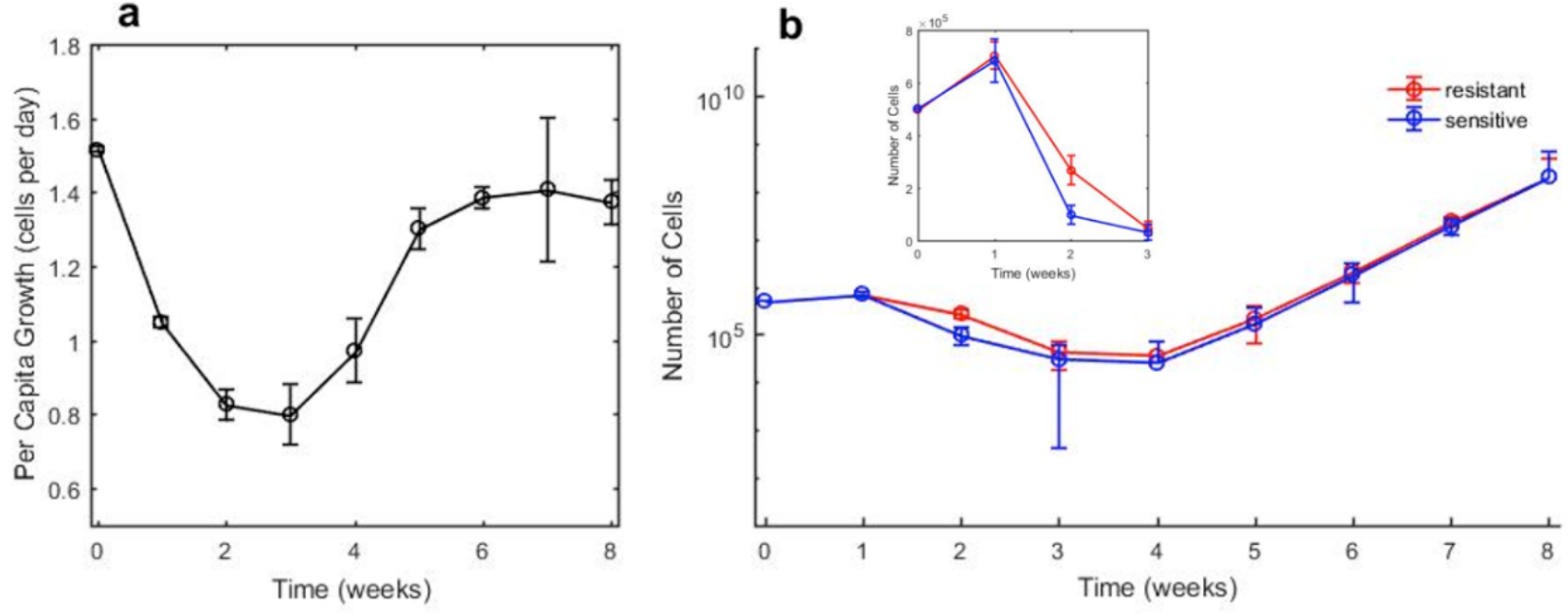
Data Driven Estimates of Phenotypic Dynamics. **a.** Per capita growth rate (birth minus death) per day of entire cell population following doxorubicin treatment **b.** Estimates of the number of resistant and sensitive cells over time are obtained by combining bulk proliferation rate with fraction estimates from model output. **Inset**: Zoom in on numeric scale of weeks 1-3 displaying higher number of resistance cells at these times.

### Model Validation Confirms Ability to Reveal Subpopulation Composition Defined by Drug Sensitivity

Without a molecular marker of drug resistance, the estimated changes in drug resistant and drug sensitive subpopulations are difficult to validate; that is, we did not know for certain that our model estimated parameters of resistant and sensitive fractions were the true estimates of the subpopulation compositions. We validated the modeling approach by generating experimental reference standards consisting of mixtures of the wild-type MCF-7 cell line with its corresponding doxorubicin resistant cell line, the MCF-7/ADR. We evaluated the ability of the model to estimate the subpopulation composition of each reference mixture. In **Figure 5a**, the two-state model fit for each mixture is overlaid on the dose-response data for each mixture. In **Figure 5b**, the measured proportion of wild-type and MCF-7/ADR cells are plotted as the line of unity against the parameter estimations of the percent of resistant cells from the model output (**Table 2**). The measured proportions of WT and ADR cells fall within the 95 percent confidence intervals on the parameter estimations of the model, with a coefficient of determination of (R-squared) value of 0.986 (**Fig 5b**).

**Figure 5.**
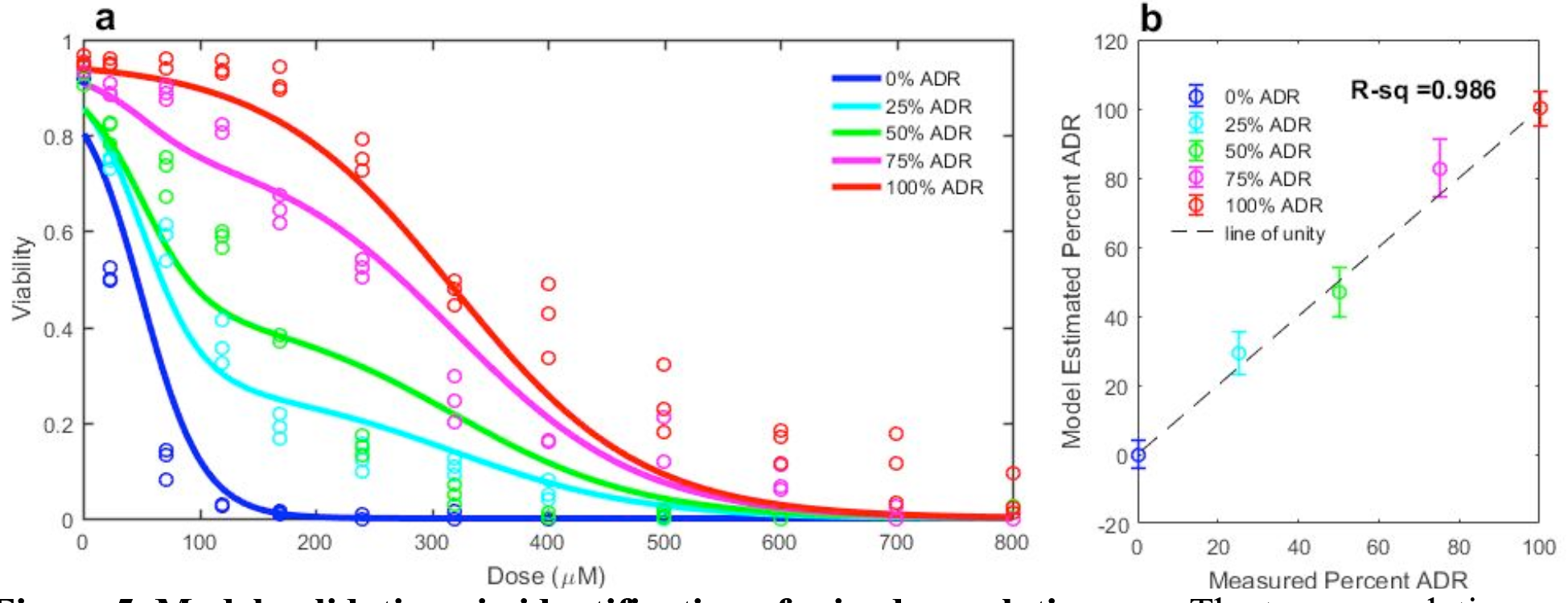
Model validation via identification of mixed populations. **a.** The two-population model fit for each mixture of MCF-7 ADR resistant cell line and WT-MCF-7 cell line drug sensitivity overlaid with the experimentally measured results. **b.** Model-estimated percent of ADR cell line in mixture versus the measured ADR cell percentage using fluorescence cell counting. The measurements fall within the 95% confidence intervals of the parameter estimations, with a coefficient of determination (R-squared) value of 0.986.

**Table 2.** Model Validation Phase Demonstrates Identifiability of Population Composition. Measured versus two population model estimated parameters indicate model’s ability to reveal the known proportions of the MCF-7 ADR resistant cell line mixed with the wild-type MCF-7 cell line. Measured LD50 values and slopes indicate the LD50 and slope of the isolated pure MCF-7 ADR cells and pure wild-type MCF-7 cells. Resistant fractions are measured precisely using the Nexcelom fluorescence imaging counter of the number of green fluorescent wild-type MCF-7 compared to the total number of brightfield cells counted.

|  | LD50<sub>ADR</sub> | slope<sub>ADR</sub> | LD50<sub>WT</sub> | slope<sub>WT</sub> | frac<sub>0%ADR</sub> | frac<sub>25%ADR</sub> | frac<sub>50%ADR</sub> | frac<sub>75%ADR</sub> | frac<sub>0%ADR</sub> |
| --- | --- | --- | --- | --- | --- | --- | --- | --- | --- |
| Measured | 374.9 | 0.0096 | 24.8 | 0.072 | 0 | 0.28 | 0.51 | 0.75 | 0.96 |
| Model | 304.2 | 0.0115 | 22.1 | 0.055 | 1.2e-5 | 0.38 | 0.53 | 0.84 | 1 |

## Discussion

The key results of this combined experimental and modeling approach are to demonstrate a time-dependency in the response to a clinically relevant pulse-dose chemotherapy treatment and to provide a modeling system that outputs estimates of the composition of a cancer cell population with functional heterogeneity in its chemoresistance. While the exact mechanism to explain the estimated resistant and sensitive compositional changes is not fully elucidated in this work, a platform for investigation of the subpopulation dynamics is the key contribution of this study. We have observed a consistent dynamic population level response in drug sensitivity following initial drug exposure, and have shown that this can be best described through a model of changing heterogeneous subpopulation compositions. We have demonstrated the ability of this modeling system to reveal population compositions defined by drug sensitivity. These subpopulations were not previously able to be identified due to the complex nature of drug resistance mechanisms. This model system is novel in that it is driven from dynamic dose-response data only, and thus we are not brute forcing the model to fit any assumed characteristics or isolated parameter measurements. This property allows the model system to be applied to a variety of cell-lines and drug conditions, as is demonstrated in the model validation portion in which the resistant subpopulation has extremely high resistance. Thus, while the results presented here are specific to the MCF-7 cell line and the initial pulsed-dose of doxorubicin, the approach can be applied to different cell types and drug indications to determine if the transient increase in the resistant subpopulation is more generally observed as a resistance response. While the fitted model parameters are likely to be dependent on a variety of factors both extrinsic and intrinsic, the key finding of this paper is to establish a foundation for describing observable resistance progression throughout time by being able to identify multiple resistant subpopulation levels previously unidentifiable by molecular markers alone. Future work will need to further investigate the molecular and cellular mechanisms that explain the observed dynamics.

Experimental *in vitro* models of resistance to cytotoxic chemotherapy typically utilize resistant cell lines developed via continuous exposure to increased drug concentration^30^. In most cases, drug resistant phenotypes are characterized by an end-point analysis following the stabilization of a resistant cell line. For the cell line of interest here, the MCF-7 breast cancer cell line, researchers have observed that the resistant phenotype is likely characterized by a multi-factorial process because of observable differences in morphology, gene expression, and DNA content between MCF-7 and MCF-7 resistant cell lines^25^. The MCF-7 resistant cell lines were on average larger, contained multiple nuclei, and were upregulated in genes involving metabolism, drug efflux, and down regulated in genes involved in DNA repair^25^. While these experimental observations provide us with key observables to identify as markers of resistance, they do not address the dynamical changes associated with resistance progression, nor are they all encompassing. To our knowledge, previous studies involving resistant cell lines have not reported time-resolved measurements of drug resistance following a clinically relevant pulsed dose chemotherapy treatment.

Mathematical modeling of heterogeneity in cancer cell populations has been investigated via multiple structural models^15–23^. Explorations into different therapy strategies such as optimal control theory have utilized the concept of resistant and sensitive cells within a tumor or cancer cell population^17–23^. Many models of *in vitro* and *in vivo* cancer progression utilize compartmental ordinary differential equations and partial differential equations. In these models, a number of key assumptions are often made. For instance, in one model of a heterogeneous tumor, it is assumed that resistance is inversely related to proliferation rate^23^. Other models assume that all sensitive cells are susceptible to the chemotherapy, and do not account for the ability of initially sensitive cells to acquire drug resistance^17^. While these models are useful in demonstrating the theoretical response to different treatment strategies under these sets of conditions, they lack in that their predictions they have yet to be fully validated experimentally due to the lack of ability to properly identify the subpopulation levels over time experimentally.

Both experimental biologists and mathematical modelers have attempted to understand the development of resistance to cytotoxic drugs in cancer treatment. For the most part, experimental biologists have discovered many useful markers of resistance, and have proposed explanations for how these markers contribute to the observed drug resistance. However, these static observations fail to reveal how resistance develops in time in a clinically relevant setting. Mathematical modelers have developed theoretical equations to predict tumor development *in vitro* and *in vivo*, but none have been fully validated by experimental observations. This work attempts to bridge the gap between these two theories of knowledge, by measuring drug resistance over time and developing a model with parameters driven exclusively by the observed results, not by separate isolated experiments. In this way, our work is unique in that it captures the dynamics of resistance development that have not been well-studied and attempts to model what is able to be measured only. This simple modeling and experimental system should allow various types of theoretical math models of resistant and sensitive subpopulations to be validated experimentally.

In this work, we investigate the dynamic changes in subpopulation composition in response to drug. In the future, time resolved subpopulation levels can be used to develop a model that describes the relative stability of drug sensitivity states and how they change in response to chemotherapy exposure. Among many questions for further investigation is the need to determine the permanence of a drug dose on the stability of drug sensitivity states. Ultimately, the results of these experimentally guided models can be used to create model systems to predict the effect of a dosing regimen on a cancer cell population over time. The goal of future studies is to use these predictions to develop and experimentally validate optimal dosing regimens to be used to combat chemoresistance.

We acknowledge the many limitations of this study. A number of key assumptions were made. We were limited in the granularity of our modeling system due to the experimental limitation of a limited dosing scheme. Because of the extreme bottle-necking of the population due to the initial pulse-treatment, we only were able to extract a small number of cells at each week for dose-response, and thus could only measure the viability reliably at 12 distinct doses at each week. For this reason, we were not able to implement a multipopulation model with more than two states due to a lack of statistical power to significantly resolve differences in subpopulation compositions in time. We acknowledge that a model of only two subpopulations does not capture all possible subpopulations, but believe that this model represents a useful simplification of resistance development. In the model validation phase, we were able to estimate the fractions of cells in each state with reasonable accuracy (**Fig 5b**), however, the estimate of LD50 of the resistant population had more than a 20 percent error from the measured LD50 of the isolated MCF-7/ADR dose response data set (**Table 2**). It is possible that discrepancies may arise from biological interactions between wild-type and resistant cell lines that actually lower the effective resistance of the resistant cell lines. A goal of future studies is to investigate the role of cell-cell interactions, along with further modeling simulations.

## Conclusion

In summary, our analysis indicates that the response to pulsed chemotherapy is time-dependent and amenable to analysis using mathematical modeling to reveal subpopulation compositions. We anticipate that some of the most exciting future advances will combine new experimental and modeling approaches to develop dynamic models of chemoresistance development.

## Acknowledgements

The authors are grateful for support from: the American Association for Cancer Research and the Breast Cancer Research Foundation through a Translational Breast Cancer Research (#14-60-26-BROC to A. Brock). We thank the National Institutes of Health for funding through NIBIB T32 EB007507, NCI 1R01CA129961, NCI 1U01CA142565, U01CA174706, and CPRIT RR160005. T.E.Y. is a CPRIT Scholar of Cancer Research.

